# First evidence of host range expansion in virophages and its potential impact on giant viruses and host cells

**DOI:** 10.1101/780841

**Authors:** Said Mougari, Nisrine Chelkha, Dehia Sahmi-Bounsiar, Fabrizio Di Pinto, Philippe Colson, Jonatas Abrahao, Bernard La Scola

**Affiliations:** Unité MEPHI, Aix-Marseille Univ., Institut de Recherche pour le Développement (IRD), Assistance Publique - Hôpitaux de Marseille (AP-HM), 19-21 boulevard Jean Moulin, 13005 Marseille, France; IHU Méditerranée Infection, 19-21 boulevard Jean Moulin, 13005 Marseille, France; Laboratório de Vírus, Departamento de Microbiologia, Instituto de Ciências Biológicas, Universidade Federal de Minas Gerais, Belo Horizonte, Minas Gerais, Brazil- postal code 31270-901

**Author notes:** Corresponding author (J.A.);. (B.L.S.).

## Abstract

Virophages are satellite-like double stranded DNA viruses whose replication requires the presence of two biological entities, a giant virus and a protist. In this report, we present the first evidence of host range expansion in a virophage. We demonstrated that the Guarani virophage was able to spontaneously expand its viral host range to replicate with two novel giant viruses that were previously nonpermissive to this virophage. We were able to characterize a potential genetic determinant of this cross-species infection. We then highlighted the relevant impact of this host adaptation on giant viruses and protists by demonstrating that coinfection with the mutant virophage abolishes giant virus production and rescues the host cell population from lysis. The results of our study help to elucidate the parasitic lifestyle of virophages and their interactions with giant viruses and protists.

## Introduction

Host range is defined as the number and nature of hosts in which a virus can multiply^1^. This parameter is predicted to play a determinant role in virus pathogenicity, maintenance in nature and epidemiology^2^. Some viruses have evolved the capacity to expand their host range by biologically adapting to novel hosts. This phenomenon is known as host range expansion and requires the selection of specific mutations that enable a given viral species to replicate in a novel host ^3^. In bacteriophages, such host range mutations target mostly the genes encoding the viral tail and baseplate involved in attachment to a receptor at the host cell surface^4,5^.

Virophages are double-stranded (ds) DNA viruses that combine features of both satellite viruses and autonomous *bona fide* viruses. On the one hand, virophages replicate only in the presence of an associated giant virus coinfecting their host cell^6,7^. This behavior is similar to the dependence of satellite viruses on their helper virus^8^. On the other hand, the level of complexity of the capsids and genomes of virophages make them closer to autonomous viruses, such as adenoviruses, than to satellite viruses^9,10^.

Studying the mechanism of host range in virophages is interesting because their replication requires the presence of two types of hosts, a giant virus and a cellular host of this giant virus (a protist). Sputnik, the first virophage to be isolated, was described in 2008^6^. Sputnik is able to infect mimiviruses of the three phylogenetic lineages (A, B and C) whose members replicate in *Acanthamoeba* spp.^6,11^. In contrast, another virophage named Zamilon was found to exhibit a narrower host range, being able to replicate with mimiviruses from lineages B and C but not with those belonging to lineage A^12^. The resistance of lineage A mimiviruses to Zamilon has been linked to the presence of a defense system^13^, named MIMIVIRE, which is composed of a helicase, a nuclease and the product of a gene containing 4 repeats identical to fragments of the Zamilon genome (called Target Repeat-Containing gene – *trcg*). Trcg has been proposed to serve as a guide to target a given genetic element for its suppression (e.g., Zamilon). Although the MIMIVIRE mechanism of action is still controversial^14^, the host range of Zamilon has been efficiently expanded to mimiviruses of lineage A by silencing the MIMIVIRE genes^13^ and, more recently, by knocking out the putative canonical gene of the system using homologous recombination^15^. The MIMIVIRE system has also been successfully tested in bacteria, restoring susceptibility to antibiotics through the transformation of bacteria with plasmids containing *trcg* targeting antibiotic-resistance genes^16^. Nevertheless, host range expansion mediated by spontaneous mutations has never been described in virophages. Moreover, to date, Sputnik, Zamilon and other virophages were almost exclusively challenged with mimiviruses from the three lineages, A-C, but not with distant mimivirus relatives. The only virophage tested with a distant mimivirus was Mavirus. Mavirus replicates only in the marine flagellate *Cafeteria roenbergensis* coinfected with a distant mimivirus named CroV^17^. Moreover, once this virus begins to replicate, it has been shown that Mavirus acts as an agent of adaptive immunity that protects its host cells from CroV infection^18,19^.

In this study, two recently isolated distant mimiviruses, known as Tupanvirus Deep Ocean and Tupanvirus Soda Lake, were challenged with virophages^20–22^. We report a preliminary identification of the first mechanism of host range expansion in virophages. We were able to characterize the genetic component involved in this process. We then conducted a comprehensive study to investigate the ecological impact of virophage host range expansion through its effect on the novel virus host as well as on the host cell population. The involvement of the mutation in host range expansion of the virophage is discussed.

## Results

### First evidence of host range expansion in virophage

We used distant mimivirus relatives ^21^ Tupanvirus Deep Ocean and Tupanvirus Soda Lake as models to study the host range of virophages and their ability to replicate in amoebae infected with other mimiviruses than those belonging to the three lineages (A, B and C) of the family *Mimiviridae*^23^. Three virophages characterized in earlier studies were assayed with these giant viruses, including Guarani, Sputnik and Zamilon virophages^6,12,24^. These virophages were previously propagated in *Acanthamoeba castellanii* cells coinfected with mimiviruses and then purified. *A. castellanii* cells were inoculated with each Tupanvirus strain at a multiplicity of infection (MOI) of 10. The same MOI was used for each virophage. The replication of each virophage was then assessed by quantitative PCR (qPCR) at 0, 24 hours (h) and 48 h postinfection (p.i.). Finally, the increase in the amount of virophage DNA was calculated using the delta Ct method considering time points 0 and 48 h p.i. Sputnik and Zamilon were able to infect and replicate with both Tupanvirus Deep Ocean and Tupanvirus Soda Lake (Fig. 1a and 1b). Unexpectedly, it was not possible to detect Guarani replication with Tupanvirus Deep Ocean or Tupanvirus Soda Lake (Fig. 1a and 1b).

**Figure 1:**
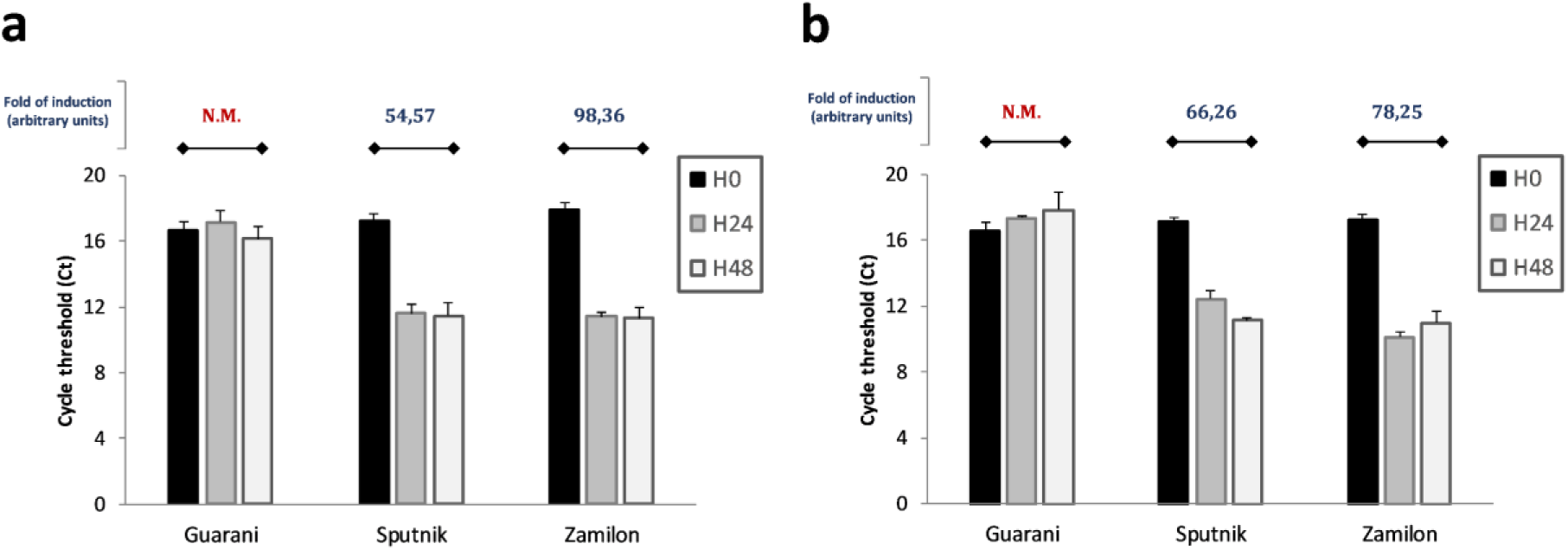
Guarani wild-type virophage is not able to replicate with Tupanviruses in primo-coculture. Histograms depicting the genome replication of Guarani, Zamilon and Sputnik in *Acanthamoeba castellanii* coinfected with Tupanvirus Deep Ocean (**a**) or Tupanvirus Soda Lake (**b**). The DNA replication of each virophage at times 0, 24 and 48 h p.i. was measured by quantitative real-time PCR. The increase in the amount of virophage DNA (fold of induction) was then calculated using the delta Ct method considering the difference between the Ct values specific to each virophage at times H0 and H48. Error bars, standard deviation. N.M.: No multiplication.

After the lysis of host cells coinfected with Guarani and Tupanvirus Deep Ocean or Tupanvirus Soda Lake, each culture supernatant was filtered through 0.22-µm-pore filters to remove giant virus particles. The obtained supernatants containing only Guarani particles were subsequently used to infect fresh *A. castellanii* cells simultaneously inoculated with Tupanvirus Deep Ocean or Tupanvirus Soda Lake. The replication of the virophage was then measured by qPCR. In these assays, Guarani remarkably infected and successfully replicated with both Tupanvirus Deep Ocean and Tupanvirus Soda Lake after only one passage with each virus (Fig. 2a and 2b). We interpreted this phenomenon as the result of a mechanism of host range expansion that allowed Guarani to replicate with new viral hosts previously resistant to this virophage. A small fraction of the Guarani population in our stocks was likely composed of an emergent mutant able to infect Tupanvirus. We found that Guarani isolated from Tupanvirus Deep Ocean culture supernatant was able to replicate with Tupanvirus Soda Lake and *vice versa*. Taken together, these results present the first evidence of a host range expansion in the virophage.

**Figure 2:**
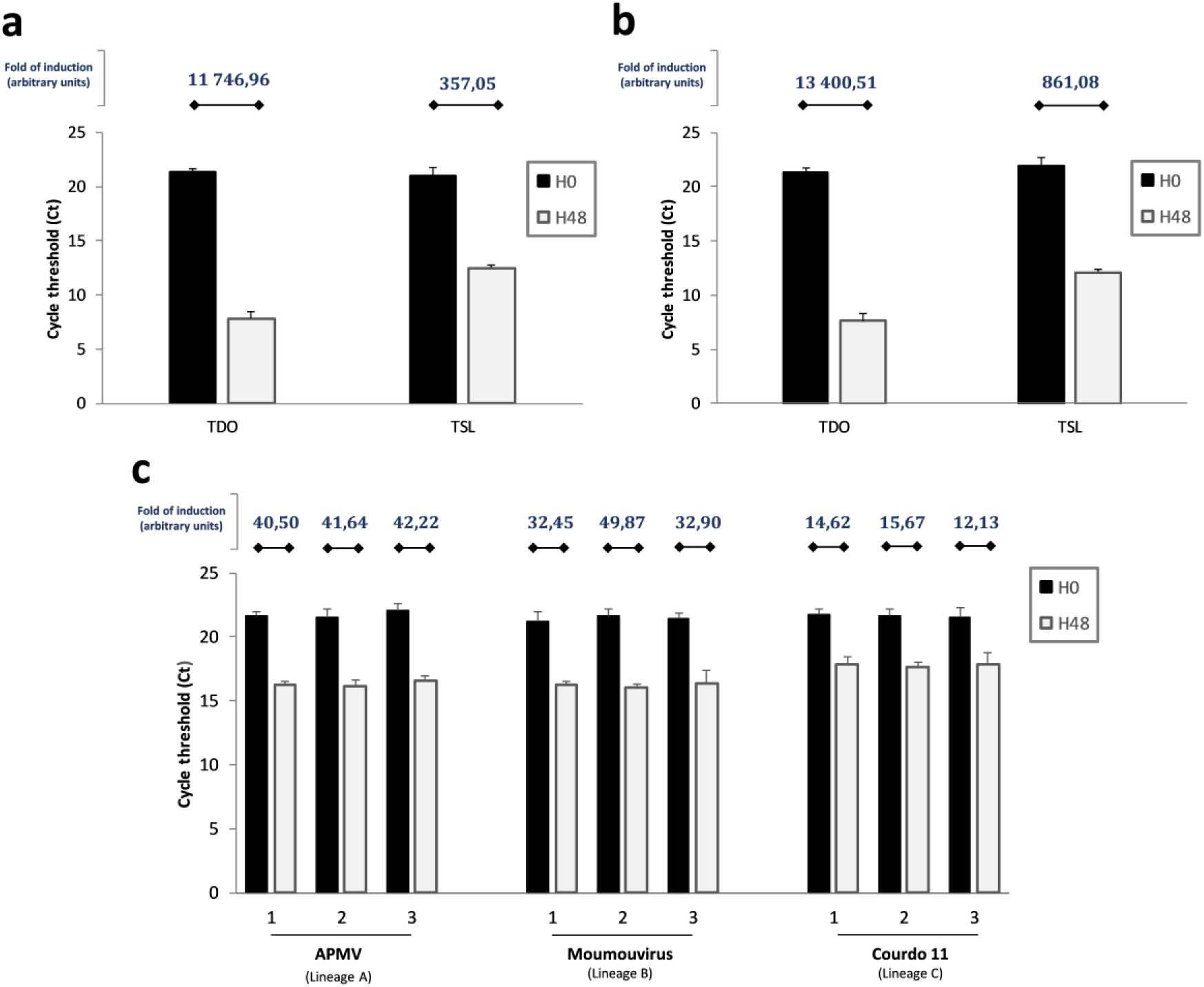
Expanded host range of Guarani isolated from Tupanviruses coculture. Graph depicting genome replication of Guarani isolated from Tupanvirus Deep Ocean supernatant (**a**) or from Tupanvirus Soda Lake supernatant (**b**) with each Tupanvirus strain. (**c**) Replication of Guarani isolated from APMV (1), Tupanvirus Deep Ocean (2) or Tupanvirus Soda Lake (3) supernatants with different mimiviruses belonging to the three phylogenetic lineages A, B and C. The DNA replication of the virophage at times 0 and 48 h p.i. was measured by quantitative real-time PCR. The increase in the amount of virophage DNA (fold of induction) was then calculated using the delta Ct method considering the difference between the Ct value specific to virophage at times H0 and H48. Error bars, standard deviation. TDO: Tupanvirus Deep Ocean. TSL: Tupanvirus Soda Lake.

We then analyzed whether host acquisition of Guarani was associated with a fitness trade-off with the prototype isolate of genus *Mimivirus*, Acanthamoeba polyphaga mimivirus (APMV), which has been used beforehand to propagate the wild-type genotype of Guarani^24,25^. Indeed, experimental evolution studies for other viruses have shown that increasing the virus fitness in one host could result in a fitness penalty in another host^1^. We compared the replication efficiency of Guarani before and after its passages with each Tupanvirus strain using APMV as a giant virus host. Figure 2c shows that even after the acquisition of new virus hosts, the virophage maintained its capacity to replicate with APMV. In addition, the replication efficiency of Guarani isolated from Tupanvirus Deep Ocean and Tupanvirus Soda Lake supernatants with APMV was similar to that of the original strain of Guarani propagated with APMV (Student’s t-test, p> 0.66 and p >0.69, respectively). Similar results were obtained with Moumouvirus and Megavirus Courdo 11, which are mimiviruses from lineages B and C, respectively^26,27^ (Fig. 2c).

### Potential host range mutation associated with the host acquisition

In bacteriophages, host range expansion involves several molecular paths. Indeed, spontaneous or induced mutations in the long tail fiber gene have enabled some phages to infect new bacterial hosts^28–32^. To identify the genetic features responsible for the expanded host range of Guarani, we sequenced the genome of this virophage obtained before and after the passage with each Tupanvirus. The Guarani genome consists of a circular dsDNA genome of 18,967 base pairs encoding 22 predicted genes very similar to Sputnik^24^. We found that the virophage isolated from APMV supernatant has no mutations in its genome compared to the original strain that we sequenced previously^24^. We therefore consider this strain to be the wild-type genotype of the virophage. Interestingly, genome analysis of Guarani isolated from Tupanvirus supernatant revealed the emergence of a new subpopulation of the virophage that shows a deletion in its genome. This deletion consists of a loss of an 81 nucleotide-long sequence located in ORF 8 that encodes a collagen-like protein (Supplementary Fig. 1, Table S1). The deletion has been detected in Guarani purified from both Tupanvirus Deep Ocean and Tupanvirus Soda Lake. We then aimed to confirm the results of the sequencing experiment by designing a PCR system that targets the deletion site, as shown in Figure 3a. We were able to confirm the presence of two genotypes of Guarani isolated from Tupanvirus supernatant versus only one genotype in APMV supernatant (Fig. 3b). The band corresponding to each genotype detected in the Tupanvirus supernatant was recovered from the 2% agarose gel. Sanger sequencing was then performed on each band and confirmed that the deletion-mutation of the 81-nucleotide-long stretch is localized in ORF 8 of the mutant virophage. The same procedure was performed with Guarani cultivated with APMV and confirmed the presence of the wild-type genotype, but the mutant was not detected in this condition.

**Figure 3:**
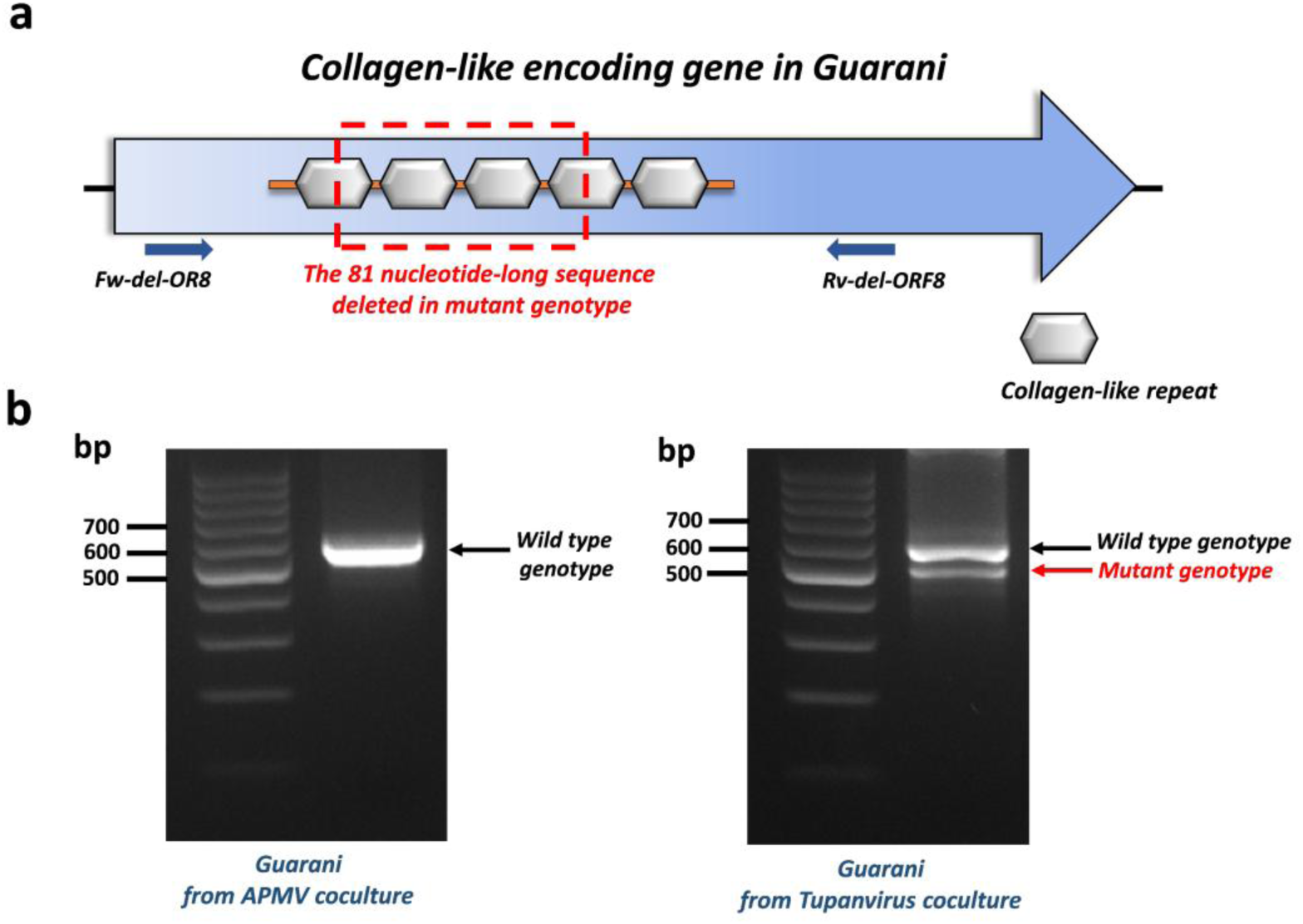
Characterization of the mutant genotype of Guarani isolated from Tupanvirus coculture. (**a**) Schematic representation of the collagen-like gene in Guarani. This gene contains 5 collagen-like repeats of 27 nucleotides each. The mutant genotype shows a deletion of an 81 nucleotides sequence that affects 4 collagen-like repeats, from which 2 repeats are completely lost. (**b**) PCR characterization of the mutant genotype detected in Tupanvirus supernatant but not in APMV coculture. The primers used for this PCR system (Fw-del-OR8 and Rv-del-OR8) target the regions located around the deletion site. The genotype corresponding to each band was then confirmed by Sanger sequencing.

### Evolution of the mutant virophage and Tupanvirus during a serial passage experiment

Virus adaptation to new hosts combines two steps: first, the introduction of mutations required to allow the virus to infect new hosts and then its ability to maintain the mutations during spread within the host population^33^. With this in mind, we investigated the capacity of Guarani to maintain the deletion during passage experiments using Tupanvirus Deep Ocean as a virus host. This Tupanvirus strain was selected because it enables the virophage to replicate with higher replication efficiency (Fig. 2a and 2b). We continuously cocultured the mutant genotype 5 times in *A. castellanii* and subsequently characterized its progeny after each passage using our PCR system targeting the deletion site (Fig. 4a). The same procedure was carried out for APMV and Guarani wild-type (control) (Fig. 4b). Passages with Tupanvirus promoted the propagation of the mutant genotype, rather than the wild-type strain (Fig. 4a). Clearly, according to the PCR product intensity, after several passages with Tupanvirus, the amount of mutant Guarani DNA evolved to decrease (Fig. 4a). Such a decrease was not observed in the control (Fig. 4b). These results are intriguing given the high replication efficiency of the mutant genotype observed with Tupanvirus Deep Ocean (Fig. 2a). At the same time, we noticed a progressive increase in the host cell population survival over the passages (data not shown). This probably raises questions regarding the virulence of the mutant genotype toward its new virus host, Tupanvirus. Indeed, it is known from our previous study that Guarani impaired the infectivity of its viral host APMV, resulting in a decrease in amoebae lysis^24^. Nevertheless, as this is the first time that we describe a host range expanding mutation in virophage, we expected the mutant genotype to have a distinct virulent profile toward its novel virus host than the wild-type strain toward APMV. To test this hypothesis, we first started by studying the evolution of Tupanvirus infectious particle production during a passage experiment in the presence of mutant Guarani. Figure 4c shows Tupanvirus titers during a ten-passage experiment in the presence of the virophage measured by end-point dilution method. Prior to end point dilution, the virus supernatant was submitted to heat treatment at 55°C for 30 min^34^. This treatment efficiently inactivated the mutant virophage without reducing the titer of viable virus particles (Supplementary Fig. 2). We found that infection with the mutant strain severely modifies the trajectory of Tupanvirus in the model. While the virus was found to increasing its titer over passages in the absence of the virophage, the presence of the latter caused a drastic decrease in virus propagation (Fig. 4c). Moreover, at the ninth passage, we were intrigued by the absence of cell lysis and any evident cytopathic effect on the host population. This observation probably indicates that the mutation not only enabled Guarani to replicate with Tupanvirus but also increased its virulence to the point of inducing the elimination of the giant virus in the model.

**Figure 4:**
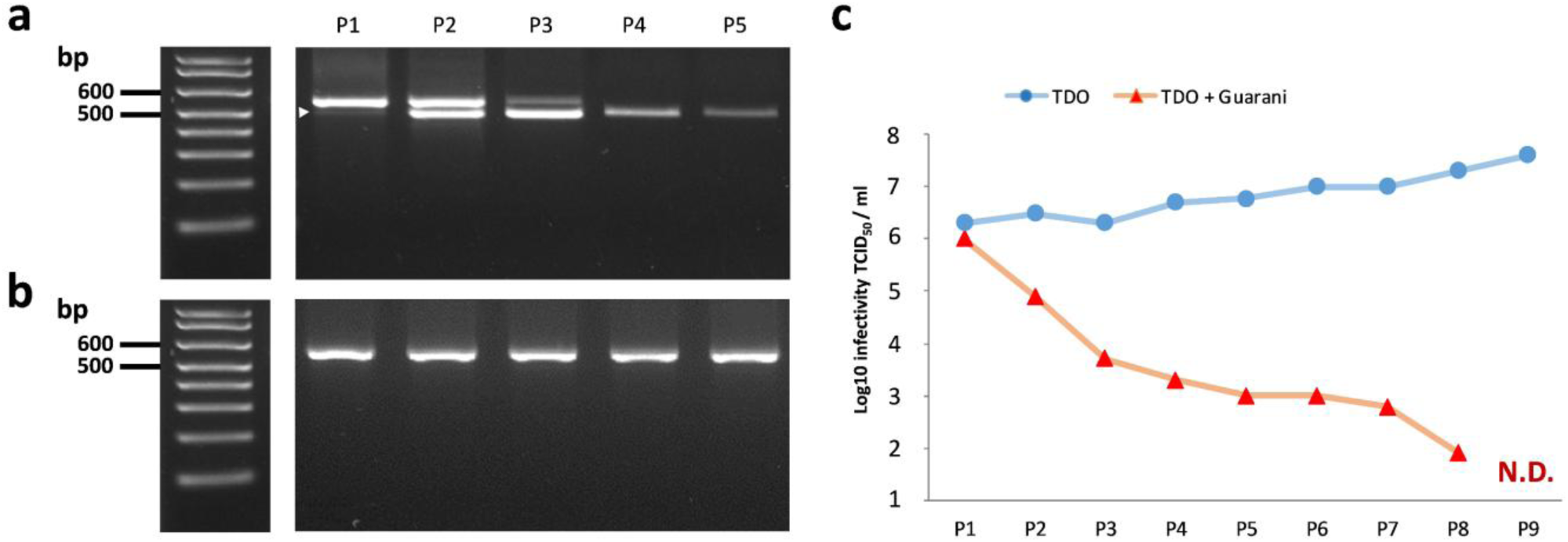
Selection of mutant Guarani during serial passage in Tupanvirus Deep Ocean. (**a**) Selection of the Guarani mutant genotype coinfecting Tupanvirus during a 5-passage experiment could be visualized by PCR (lower band-arrow). (**b**) The same experiment was repeated using APMV (control), revealing the selection and maintenance of wild-type Guarani through passages. (**c**) Tupanvirus titer during 10 passages in the presence and absence of mutant virophage measured by the end point method. The results with the wild-type genotype were similar to those obtained in the absence of virophage (not shown). P: Passage. TDO: Tupanvirus Deep Ocean. N.D.: Not detectable.

### Virophage infection leads to Tupanvirus elimination

To go further into this story and investigate the effect of mutant Guarani on the replication of Tupanvirus during a one-step growth curve, *A. castellanii* cells were coinfected with Tupanvirus Deep Ocean and Guarani mixture, containing both wild-type and mutant genotype (according to PCR), at MOIs of 10. The Guarani used in this study was isolated and then purified from the second passage with Tupanvirus in which the mutation was detected. We first quantified the impact of virophage on genome replication of Tupanvirus at 48 h p.i. Figure 5a shows that mutant Guarani decreases the replication of Tupanvirus DNA by approximately 3-fold. This low inhibition rate appears relatively similar to what has been found between the wild-type strain and APMV^24^. Experimental studies supported by bioinformatic analyses have shown that in contrast to their virus hosts, virophages become active during a late phase of giant virus infection^17,24,35^. Since genome replication is an early step in the cycle of giant viruses, this property may allow them to anticipate virophage parasitism and produce some genomic copies. We then tested the effect on the production of viable particles by quantifying the titer of Tupanvirus from 0 h to 72 h p.i. Our results reveal that mutant Guarani induces a severe decrease in the production of Tupanvirus virions during the virus cycle (Fig. 5b). We found that in the absence of virophage, the virus was able to increase its titer by up to 500-fold during a one-step growth curve in amoebae. In contrast, infection with the mutant genotype prevented any increase in the virus titer. This inhibition is significantly far higher than that of the wild-type toward APMV^24^, but at 48 h p.i., we still were able to detect the production of infectious virions by end point dilution.

**Figure 5:**
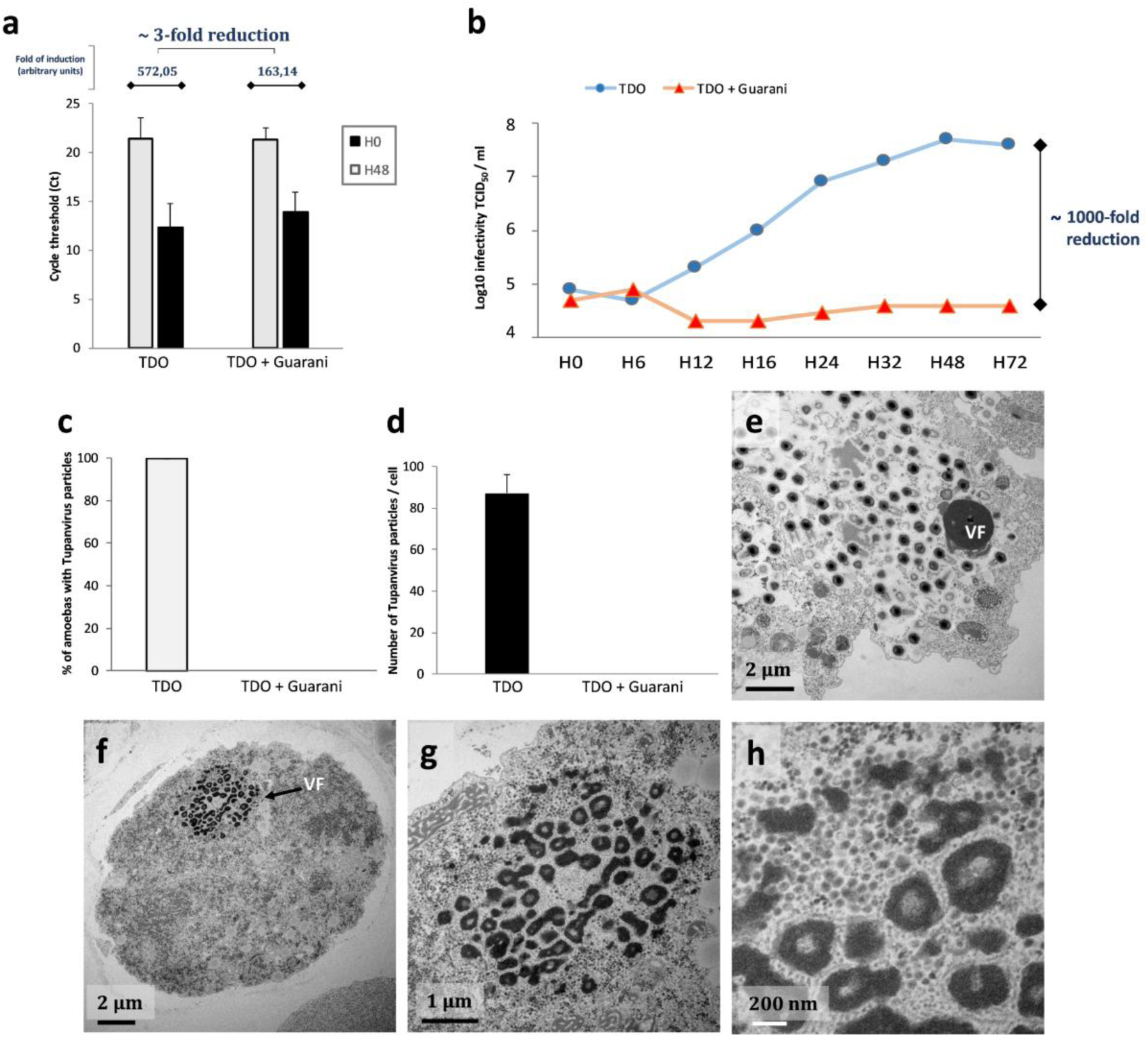
Characterization of the biological activity of mutant Guarani on Tupanvirus Deep Ocean propagation. (**a**) Genome replication of Tupanvirus in the presence and absence of mutant virophage measured by quantitative real-time PCR. The Delta Ct method was used to analyze the increase in virus DNA for each condition between times H0 and H48 as described above. Error bars, standard deviation. (**b**) A one-step growth curve of Tupanvirus in the presence and absence of mutant Guarani showing that the virophage drastically reduces the ability of the virus to produce viable virions in coinfected amoebas. (**e-h**) Transmission electron microscopy analyses show that mutant virophage induces a total inhibition of Tupanvirus morphogenesis. Graphs **c** and **d** were generated by analyzing the viral factories of 200 coinfected cells. Error bars, standard deviation. (**c**) Percentage of infected amoebas in which Tupanvirus virions have been observed in the presence and absence of mutant virophage. (**d**) The average number of Tupanvirus particles observed in each infected cell in the presence and absence of mutant virophage. (**e-h**) Transmission electron microscopy images of *A. castellanii* cells infected with Tupanvirus at 24 h p.i. (**e**) In the absence of virophage infection, the cell cytoplasm is fulfilled by mature viral particles. (**f-g**) The presence of mutant Guarani completely interrupts the production of Tupanvirus virions in coinfected cells, in which only the virophage progeny could be observed (**h**). TDO: Tupanvirus Deep Ocean. VF: Virus factory.

To gain more insight into the virophage–giant virus interaction, amoebas were coinfected with Tupanvirus Deep Ocean and mutant Guarani at MOIs of 10 and prepared for transmission electron microscopy (TEM). Remarkably, TEM images revealed that infection with mutant Guarani induces a total inhibition of Tupanvirus particle morphogenesis (Fig. 5c-5h). We scanned more than two hundred cells and quantified the number of amoebas, in which both virophage and giant virus progeny were observed (Fig. 5c-5d). Our results confirmed the absence of any simultaneous occurrence of Guarani and Tupanvirus virion production in all coinfected cells. The presence of the virophage was automatically associated with the absence of Tupanvirus virions (Fig. 5e-5h). To characterize this phenomenon, we observed virus factories infected with the mutant virophage at serial stages of coinfection (Fig. 6). Figure 6a shows that at 16 h p.i., the virus was able to produce mature virus factories, even in the presence of virophage. The first virophage progeny was observed at this step. Later in the cycle, the replication of the virophage increased progressively and caused remarkable damage to virus factories (Fig. 6b-6e). The latter appeared disintegrated, and virophage progeny were actively emerging from each of their pieces. Virophage multiplication was clearly correlated with a progressive degradation of Tupanvirus viral factories. Isolation and characterization of these structures have previously shown that they are composed of an arsenal of virus-encoded proteins involved in virus replication and assembly^36^. Therefore, there is some overlap between our observations and previous studies reporting that virophages hijack their giant virus-encoded machinery to express and probably replicate their genomes^35,37^. In this study, we clearly observed that these small viruses obtain essential elements from the factories of their host viruses to propagate.

**Figure 6:**
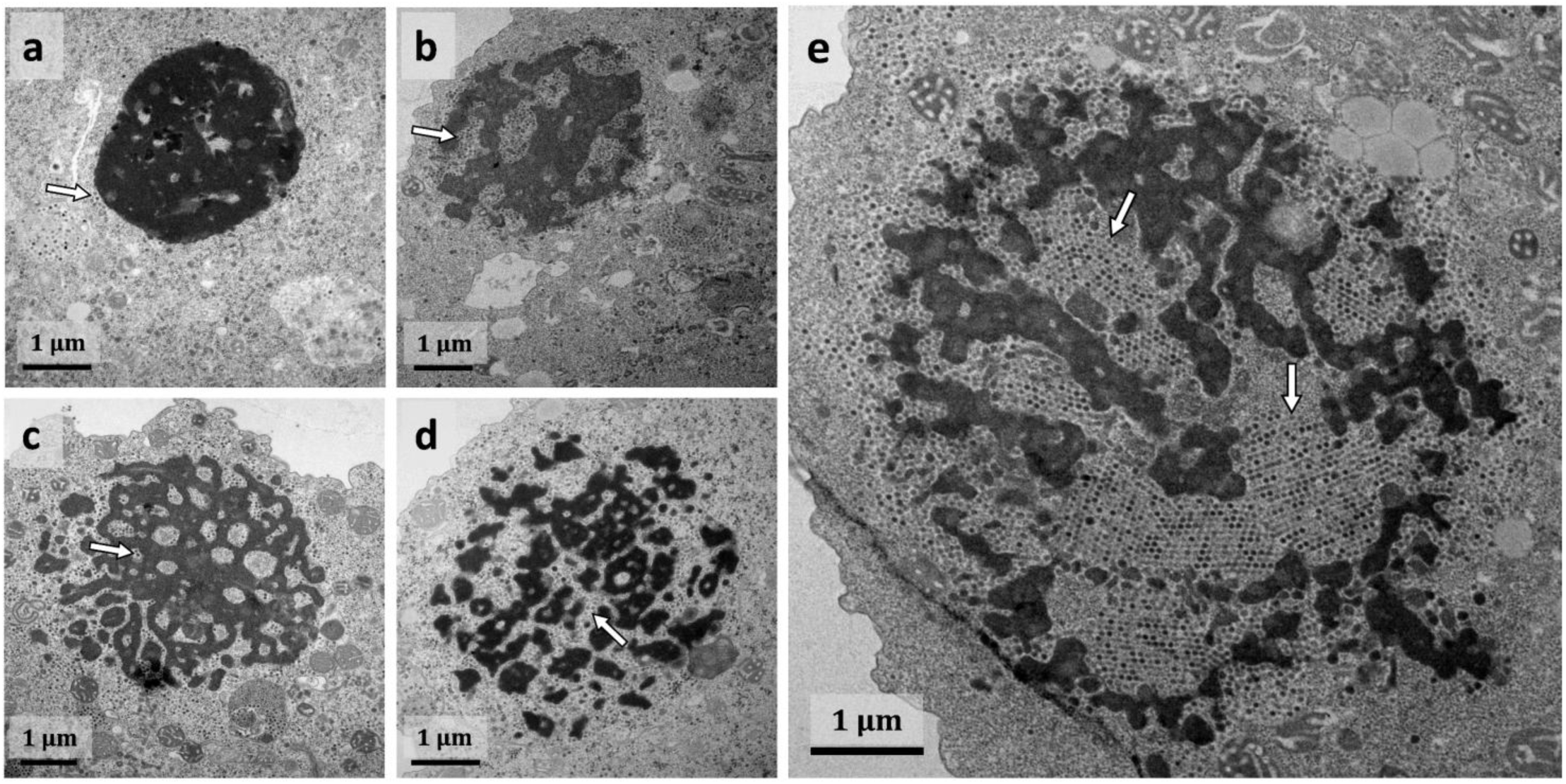
Virophage replication is associated with a progressive degradation of Tupanvirus viral factories. Transmission electron microscopy images of Tupanvirus viral factories infected with mutant Guarani at different stages of infection. (**a**) The first virophage progeny could be observed at 16 h p.i. (arrow), at this stage, the virus factories seem to be still intact. (**b-e**) Later in the cycle, the replication of the virophage (arrows) causes a remarkable fragmentation of the virus factories and seems to be correlated with their progressive degradation, affecting Tupanvirus particle morphogenesis.

However, it is still not clear how the virus was able to produce infectious virions at the end of its cycle (Fig. 5b), despite the presence of the virophage that is supposed to induce its elimination. We therefore quantified the rate of infected cells for each virus (Tupanvirus and Guarani). We found that all observed cells were successfully infected with Tupanvirus (100%). Although no virions have been produced, the presence of their volcanic-like virus factory in the cytoplasm of a given host cell was considered a sign of infection with the virus. On the other hand, virophage virions were observed in approximately 76% of host cells (152 cells from 200 cells observed). This finding means that up to 24% of amoebas were successfully infected only by Tupanvirus alone without the virophage. Two plausible explanations could justify this observation. First, the virophage suspension used here contained both mutant and wild-type genotypes; thus, the latter was not able to replicate in cells it infected. Second, the titration of virophage was carried out using qPCR which, in contrast to end-point dilution, targets both infectious and defective particles. Based on these results, we speculate that giant virus particles produced during coinfection experiments with Tupanvirus and mutant Guarani at MOIs of 10 were released from amoebas infected only by the giant virus. We therefore reduced the MOI of Tupanvirus from 0.01 to 1 and performed a dose-response experiment to virophage (Fig. 7). To improve the infection rate with virophage prior to coinfection, the virophage was incubated with Tupanvirus for 30 min at 30 °C to allow the formation of giant virus-virophage-composite. Indeed, Sputnik-like virophages are thought to enter their host cell simultaneously with their associated giant virus, attaching to its capsid fibrils ^38,39^. Figure 7a demonstrates that regardless of the MOI of Tupanvirus, higher virophage concentrations systematically reduced the titer of Tupanvirus to undetectable levels. In parallel with this result, Zamilon was not able to cause such inhibition for Tupanvirus (Fig. 7b), and simultaneous production of virophage and giant virus progenies was observed in coinfected cells (Supplementary Fig. 3). On the other hand, although mutant Guarani has caused a decrease in APMV titer, increasing MOI of this virophage was not able to completely prevent the propagation of the virus. These results confirm that Tupanvirus was unable to establish a productive infection in host cells that are simultaneously coinfected by the mutant genotype of Guarani.

**Figure 7:**
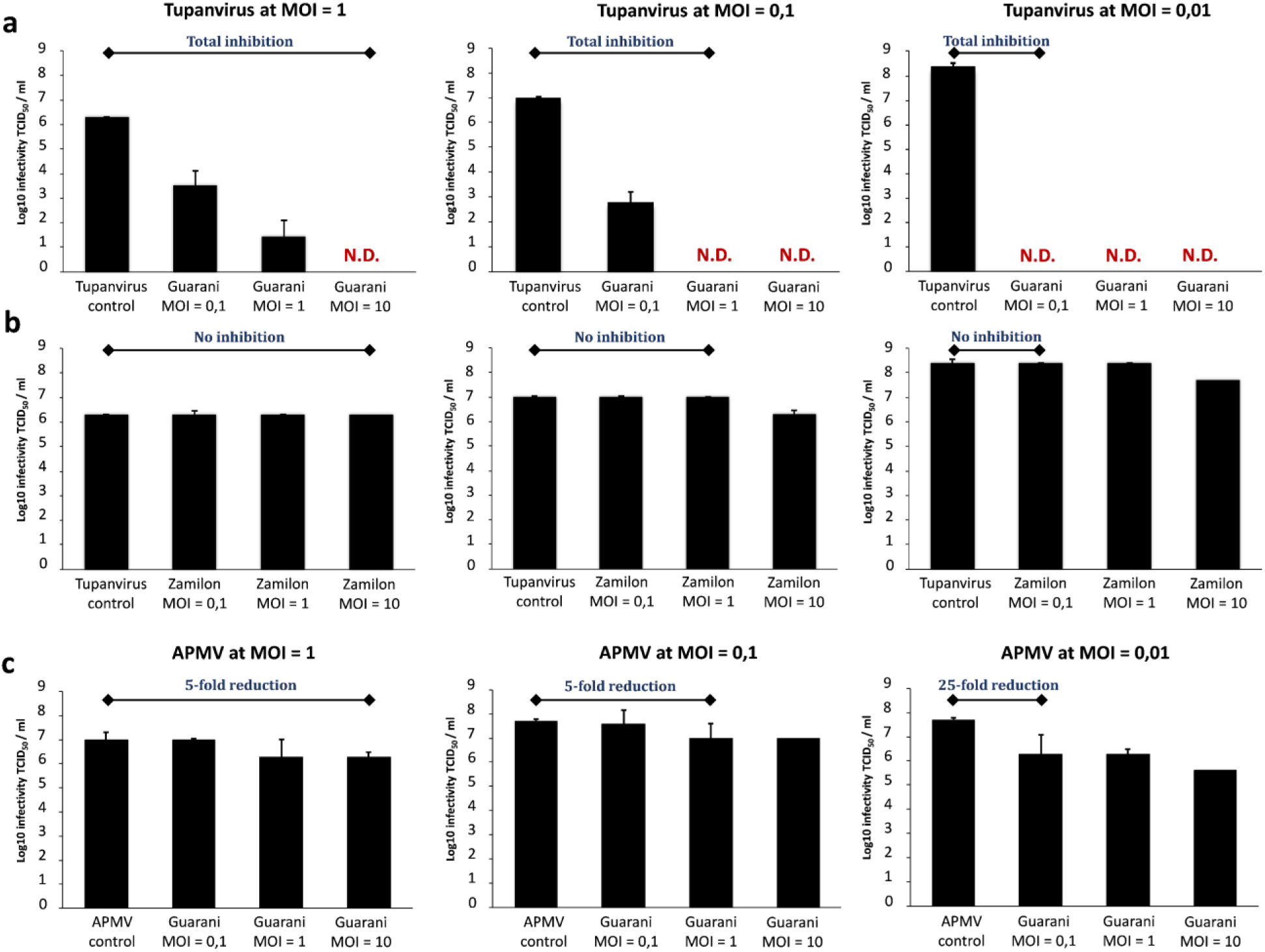
Dose response of Tupanvirus Deep Ocean to virophage. *A. castellanii* cells were coinfected with Tupanvirus at different MOIs (1, 0.1 and 0.01) and increasing MOIs (0.1, 1 and 10) of mutant Guarani. The Tupanvirus titer in the supernatant was then measured by end point dilution after complete lysis of cells or 5 days p.i. in the absence of cell lysis. (**a**) Higher concentrations of virophage cause a systematic fall in Tupanvirus titer to undetectable levels. We interpreted these observations as the results of a phenomenon of elimination induced by virophage. This phenomenon was not observed when Tupanvirus was challenged with Zamilon (**b**) nor when APMV was challenged with the mutant Guarani (**c**). Error bars, standard deviation. N.D.: Not detectable.

### Host cell population survives Tupanvirus infection in the presence of the mutant virophage

The next step was to investigate how the mutant Guarani could manipulate the stability of the host cell population. To this end, *A. castellanii* cells in PYG (Peptone, Yeast extract, Glucose) medium were simultaneously inoculated with Tupanvirus at several MOIs (from 0.01 to 1) and mutant Guarani at high MOI (MOI= 10). The concentration of the host cell population was then monitored by microscopy count from 0 h to 96 h p.i. In the absence of Guarani, Tupanvirus infection led to a dynamic decrease in cell density until causing a total cell lysis at 96 h approximatively (Fig. 8). Interestingly, at low MOIs of Tupanvirus (0.1 and 0.01), coinfection with Guarani was able to stop the propagation of the virus and thus rescued the host cell population from lysis (Fig. 8). Even in the presence of the virophage, we observed a cytopathic effect (rounded cells) and detected cell lysis with Tupanvirus at an MOI of 1. This finding may be observed because infection by Tupanvirus inevitably causes cell lysis, even when no virions could be produced. This observation is similar to what has been described for CroV^18^. However, by reducing the virus MOI, we observed that neighboring cells did not seem to be affected after lysis of infected cells, most likely because the virophage was able to efficiently neutralize the virus in coinfected cells.

**Figure 8:**
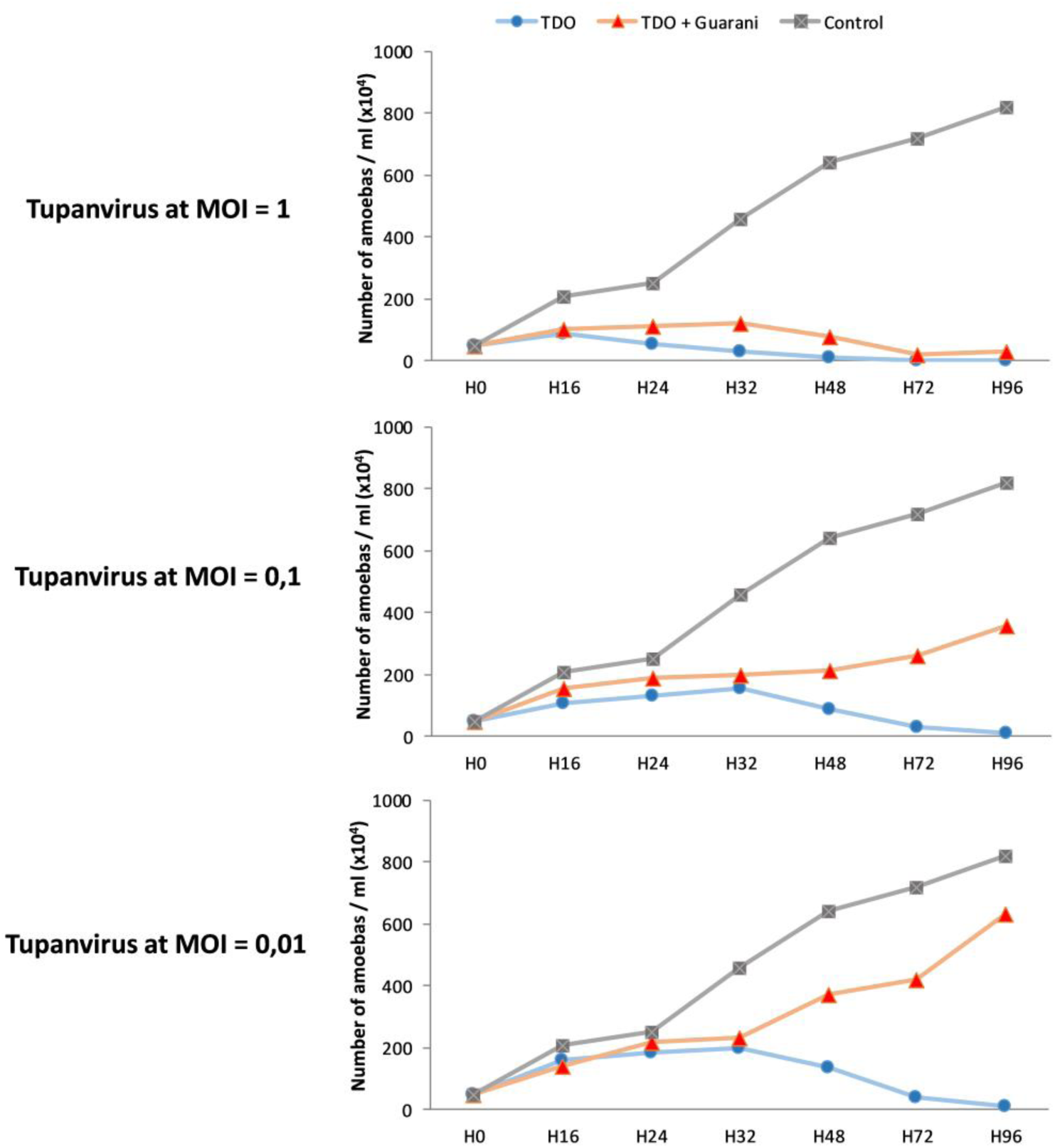
Infection with the virophage protects the host cell (amoebas) population from Tupanvirus infection. *A. castellanii* cells in PYG medium were coinfected with Tupanvirus Deep Ocean at different MOIs (1, 0.1 and 0.01) and mutant Guarani at an MOI of 10. Cell densities were then monitored by microscopy count from 0 h to 96 h p.i. Uninfected amoebas were used as controls. In the absence of virophage, regardless of Tupanvirus MOI, the virus lyses the entire host cell population at 96 h p.i. approximatively. The presence of virophage dramatically modifies the lysis of amoebas at low MOIs of the Tupanvirus. The virophage protects a portion of host cells from lysis at MOIs of 0.1 and 0.01 of Tupanvirus. At an MOI of 0.01 of Tupanvirus, the phenotype of the amoebae population seems similar to that of uninfected cells. TDO: Tupanvirus Deep Ocean.

### Tupanvirus-induced host-ribosomal shutdown prevents virophage infection

Tupanvirus is one of the most complex viruses isolated to date^22^. In addition to the distinct genetic and structural characteristics of this virus, it exhibits a remarkable biological feature never previously observed in other giant viruses. Tupanvirus is able to trigger a cytotoxic profile associated with a shutdown of ribosomal RNA (rRNA) in host and nonhost organisms^20^. This profile seems to be independent of virus replication and probably allows the virus to modulate nonhost predator organisms to increase viral survival chances in nature. In host organisms, such as *A. castellanii,* this phenomenon is mainly observed at high MOIs (MOI= 100)^20^. In this study, we investigated how this cytotoxicity could modulate virophage parasitism and notably infection by the mutant genotype of Guarani. To the best of our knowledge, such a study has never been conducted. *A. castellanii* cells were coinfected with Tupanvirus at an MOI of 100 and each virophage (Sputnik, Zamilon or mutant Guarani) at MOIs of 10. Moumouvirus, a lineage B mimivirus, was used as a control because it allows the replication of all virophages used in this study. First, at 9 h p.i., 5×10^5^ cells were collected to check the impact of Guarani on ribosomal shutdown. We found that in contrast to the controls, irrespective of mutant Guarani infection, Tupanvirus was able to induce a severe shutdown in host ribosomal RNA abundance (Fig. 9a). We then measured the replication of each virophage by qPCR and calculated their DNA amount increase using the delta Ct method considering times 0 and 48 p.i. All virophages were able to replicate with Moumouvirus at the MOI of 100 (Fig. 9b). In contrast, all of the virophages, including Guarani, failed to infect Tupanvirus at high MOI (Fig. 9c). These results suggest that the cytotoxic profile of Tupanvirus allowed it to escape virophage infection and, notably, the eradication that could be caused by Guarani.

**Figure 9:**
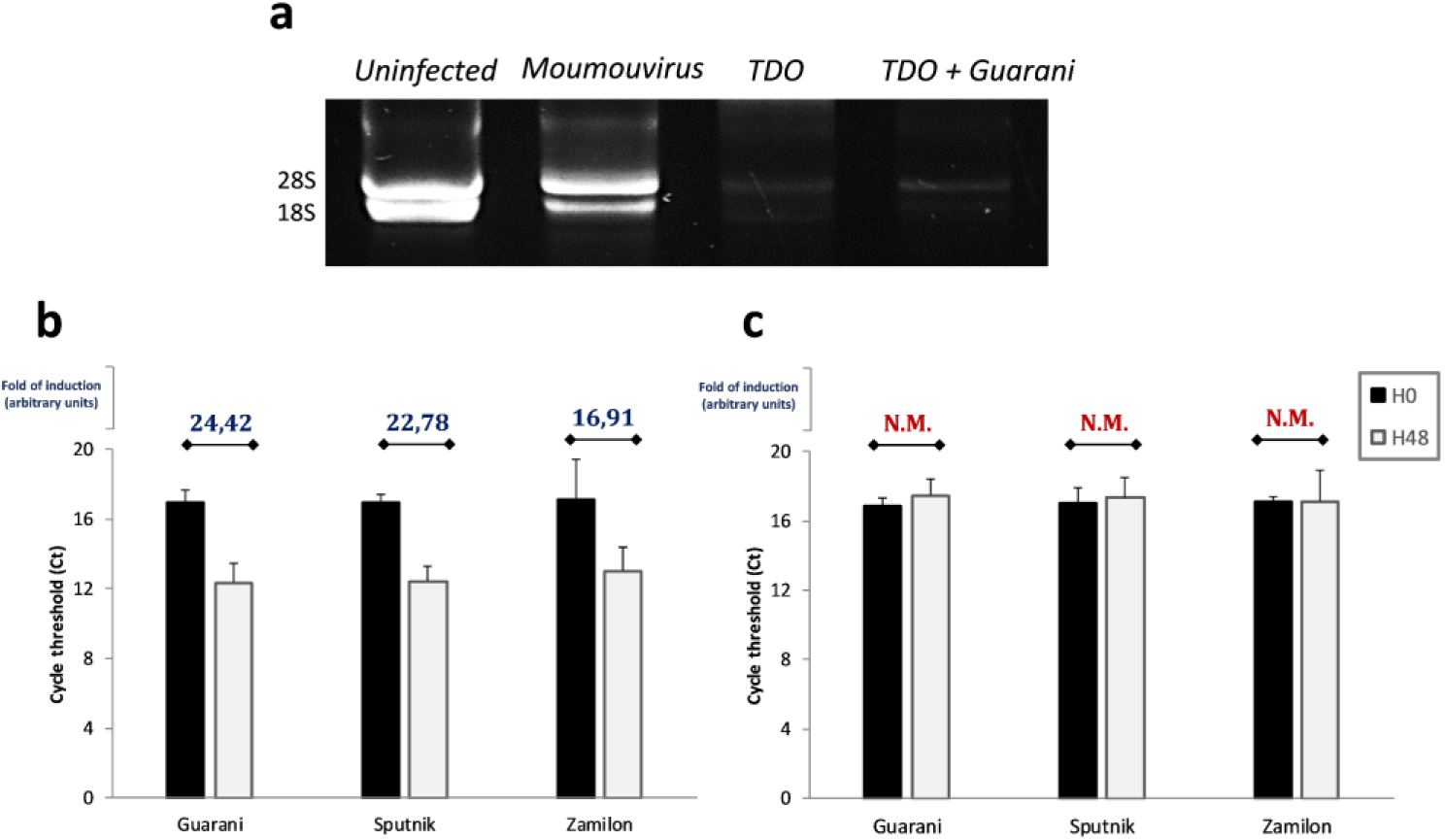
Virophage infection and rRNA shutdown induced by Tupanvirus. *A. castellanii* cells were coinfected with Tupanvirus Deep Ocean at an MOI of 100 and each virophage. (**a**) Electrophoresis gel showing the ribosomal RNA profile (18S and 28S) from *A. castellanii* in the presence and absence of Guarani. Moumouvirus was used as a giant virus control. Tupanvirus has remarkably induced a severe shutdown in rRNA even in the presence of virophage. (**b**) All virophages replicate with Moumouvirus at an MOI of 100. (**c**) All virophages failed to replicate in amoebas coinfected with Tupanvirus at an MOI of 100. Error bars, standard deviation. TDO: Tupanvirus Deep Ocean.

## Discussion

Evolution is an inevitable path for living organisms to adapt to changes in their ecosystem and to explore new environmental niches. In microorganisms, the evolutionary process has been studied in considerable detail. The process seems to be driven by two major factors: genome plasticity and selection^40^. Because of their fast replication and large population sizes, viruses offer an experimental advantage to study the evolutionary process at the laboratory^41^. Although much is known about the process of evolution in bacteriophages and many emerging zoonotic viruses and how they adapt to new hosts, this mechanism is still unclear for virophages and satellite viruses.

In this study, we report the first description of a host range expansion in virophages. We demonstrated that the Guarani virophage was able to spontaneously expand its host range to infect two novel giant viruses from the genus Tupanvirus that were initially resistant to this virophage. Host acquisition was associated with the emergence of a new genotype of Guarani that showed deletion of an 81-nucleotide long sequence from ORF8 that encodes a collagen repeat-containing protein. We found that despite this deletion, the virophage was still able to replicate with its previous virus hosts without any fitness change (Fig. 3c). One might suppose that this replication has been observed because of the presence of a background of wild-type virophage. However, almost pure mutant Guarani (isolated from passage 5 with Tupanvirus as shown in Fig. 4a) displayed a similar replication profile with mimiviruses group A, B and C, as the wild-type strain propagated with APMV (data not shown). Moreover, we have no evidence regarding the viability of wild-type virions present in the virophage after coinfection with Tupanvirus. We also demonstrated that repeated passages in the presence of Tupanvirus promoted propagation of the mutant genotype. These results provide evidence that this deletion mutation was the primary, but possibly not exclusive, determinant of host range expansion of Guarani. The first question concerns the origin of this deletion. While both Illumina sequencing and PCR failed to detect the mutant genotype in the initial isolate of the virophage, several studies have shown that host range mutations usually exist in the viral population before contact with the new host as part of the virus’s genetic diversity^3,42^. Therefore, it is possible that the mutant genotype was already present in the Guarani population that seemed not able to replicate with Tupanvirus. However, its concentration was probably under the limit of detection of our systems. Then, Tupanvirus allowed the selection of the mutant in coinfected cells. The presence of a very low amount of the mutant virophage after the first passage with Tupanvirus probably supports this hypothesis (Fig. 4a, arrow). This finding also suggests that the virophage has replicated even in the primo coinfection with Tupanvirus at levels too low to be detected by real-time PCR but high enough to expand the virophage population, enabling its multiplication in subcultures.

Several studies have reported the occurrence of spontaneous large genomic deletions in dsDNA viruses, including the giant Mimivirus, poxviruses, African swine fever virus (ASFV), and chlorella viruses^43–46^. In all these viruses, the deletion concerned sequences of several kilobase pairs (kbp) in length located around each end of the virus genomes. The ends of the genomes of these viruses seem to be highly recombinogenic in contrast to the central region, which is more subjected to selective pressures because it contains conserved core genes^43^. For ASFV, the deletion was associated with an enhancement in the capacity of the virus to replicate in Vero cells^45^. For Mimivirus, the loss of genes affected genes encoding fibers and ankyrin repeat-containing proteins. This genetic variation has given rise to a new subpopulation of viruses lacking surface fibers that were resistant to virophage^43^. Spontaneous in-frame deletions have also been reported for several other viruses, including influenza virus, photosynthetic cyanophages and severe acute respiratory syndrome coronavirus (SARS-CoV). Depending on the affected genes, these mutations could have a beneficial or deleterious impact on the viral fitness^47–49^.

In our case, the deletion affected a part of a gene. ORF 8 in Guarani is 933 base pairs in length and contains 310 amino acids^24^. The same gene was described in all Sputnik strains (with 100% amino acid identity) and was predicted to be involved in protein-protein interactions with giant viruses within factories^6,11^. This gene shows a remarkable repetitive pattern. Five repeats of 27 nucleotides could be detected. The deletion affects 4 repeats, of which 2 repeats are completely lost (Fig. 3a). Our PCR system targeting the deletion site confirmed the presence of a new subpopulation in Sputnik that shows the same deletion as Guarani even before contact with Tupanvirus. This finding probably explains the capacity of Sputnik to replicate with this virus in primo coculture (Supplementary Fig. 4).

Although it is difficult to predict the exact function that this gene could play during virophage infection, one plausible scenario is that ORF8 is implicated in the recognition and attachment to Tupanvirus proteins, probably via its multiple collagen repeats. Indeed, collagen-like motifs play a potential role in the attachment of phages to target bacteria. These motifs have been described in tail fiber proteins of several bacteriophages, and it is known from previous studies that mutations in this region are a crucial step in determining the acquisition of new hosts for these viruses^50,51^. On the other hand, collagen-like repeats have also been suggested to represent recombination hotspots for bacteriophages^50^. Interestingly, a spontaneous deletion of a region containing a collagen-like repeat has been described for the temperate *Streptococcus thermophilus* phage phi SFi21 after serial passage in bacteria. The mutant phage was remarkably unable to lysogenize its host cells^52^. It is tempting to link these observations to the study of Desnues et al^53^. The authors found that in Sputnik 2, the collagen-like gene also plays a role in recombination spot that allows the virophage to integrate itself into the Lentillevirus genome as provirophage. Therefore, the deletion in ORF8 could also involve a process of integrating Guarani into the genome of Tupanvirus that could allow the virophage to replicate. However, an experimental setting to confirm such a hypothesis is challenging and is not currently possible to set up. However, these data allow us to propose that the collagen-like gene contributes to the flexible gene content of virophages, giving them an advantage in their host-parasite interaction with giant viruses.

The second major finding of our study is that adapted virophage has not only acquired the capacity to replicate with Tupanvirus but has become highly virulent enough to induce the elimination of its novel associated virus. This observation reminds what has been described for the *S. thermophilus* phage phi Sfi21, for which a spontaneous deletion of a collagen-like repeat containing-region has transformed this temperate phage to a pure lytic phage^52^. We observed in this study that mutant Guarani did not severely affect Tupanvirus genome replication, but it seems to abolish its morphogenesis as no virions have been observed in coinfected cells. The virophage showed a clear selective inhibition toward Tupanvirus. However, the unique virophage control used in our experiment was Zamilon, and according to a previous study, this virophage does not seem to be virulent for its associated giant virus^12^. Hence, an alternative scenario is that the eradication of Tupanvirus was caused by its hypersensitivity to virulent virophage; thus, any other virulent virophage might provoke the same effect. Otherwise, apart from Guarani, the only known virulent acanthamoeba virophage is Sputnik. However, Sputnik could probably not be considered as an appropriate control because a subpopulation carrying the same mutation (as Guarani) was detected in this virophage.

We demonstrated here that mutant Guarani confers total protection to neighboring cells by abolishing the production of Tupanvirus virions in coinfected cells. This phenomenon was not observed when other mimiviruses were challenged with the mutant virophage. This finding, as well as the findings of previous studies, supports the hypothesis of a protective role of virophages toward their host cells. Indeed, Fischer *et al.* found that the virophage Mavirus has the ability to integrate its genome into that of *C. roenbergensis* cells and remains latent. Superinfection with the giant CroV triggers the expression of the provirophage, enabling virophage replication. Mavirus then acts as an efficient inhibitor of CroV by preventing its spread in neighboring cells^18^. The main advantage of integrating virophages is probably that the host cell carries a permanent antiviral weapon in its genome. However, random integration of foreign genetic elements might have considerable impacts on the cell. These repercussions might range from altering its survival to affecting its evolutionary course^54^. In this context, infection with highly virulent virophages, such as mutant Guarani, may be more advantageous for amoebae than integration of Mavirus for *C. roenbergensis*. However, the fate of the virophage is also different between these two varieties of tripartite interactions. While the Mavirus genome is efficiently preserved within the *C. roenbergensis* genome, extinction of their associated giant virus will most likely cause the extinction of highly virulent nonintegrating virophages. We observed here that mutant virophage virions were no longer viable after two months of incubation at room temperature (25 °C) and five months at 4 °C, approximatively (Supplementary Fig. 5). This finding suggests that the protection conferred to amoebae in their original habitats is probably temporary and remains dependent on viability of the virophage. Overall, this observation and that of Fischer et al., with integration of Mavirus (the predator of the predator) to *C. roenbergensis* (the prey), shows that interrelations in nature between these microorganisms are more complex than the extended Lotka-Volterra model of host–Organic Lake phycodnavirus (OLPV)–Organic Lake virophage (OLV) population dynamics proposed by Yau et al^55^.

Our results also indicate that the only way for Tupanvirus to survive virophage infection is to trigger its cytotoxic profile. This effect requires a high MOI for host organisms, and we have no evidence about the existence of such MOIs in nature. Overall, this observation and the data presented above allows us to present a new model of giant virus-virophage interaction (Fig. 10) in which the same giant virus could behave differently to virophage infection according to different parameters related to the standing genetic diversity of virophages but also to its concentration in the ecosystem.

**Figure 10:**
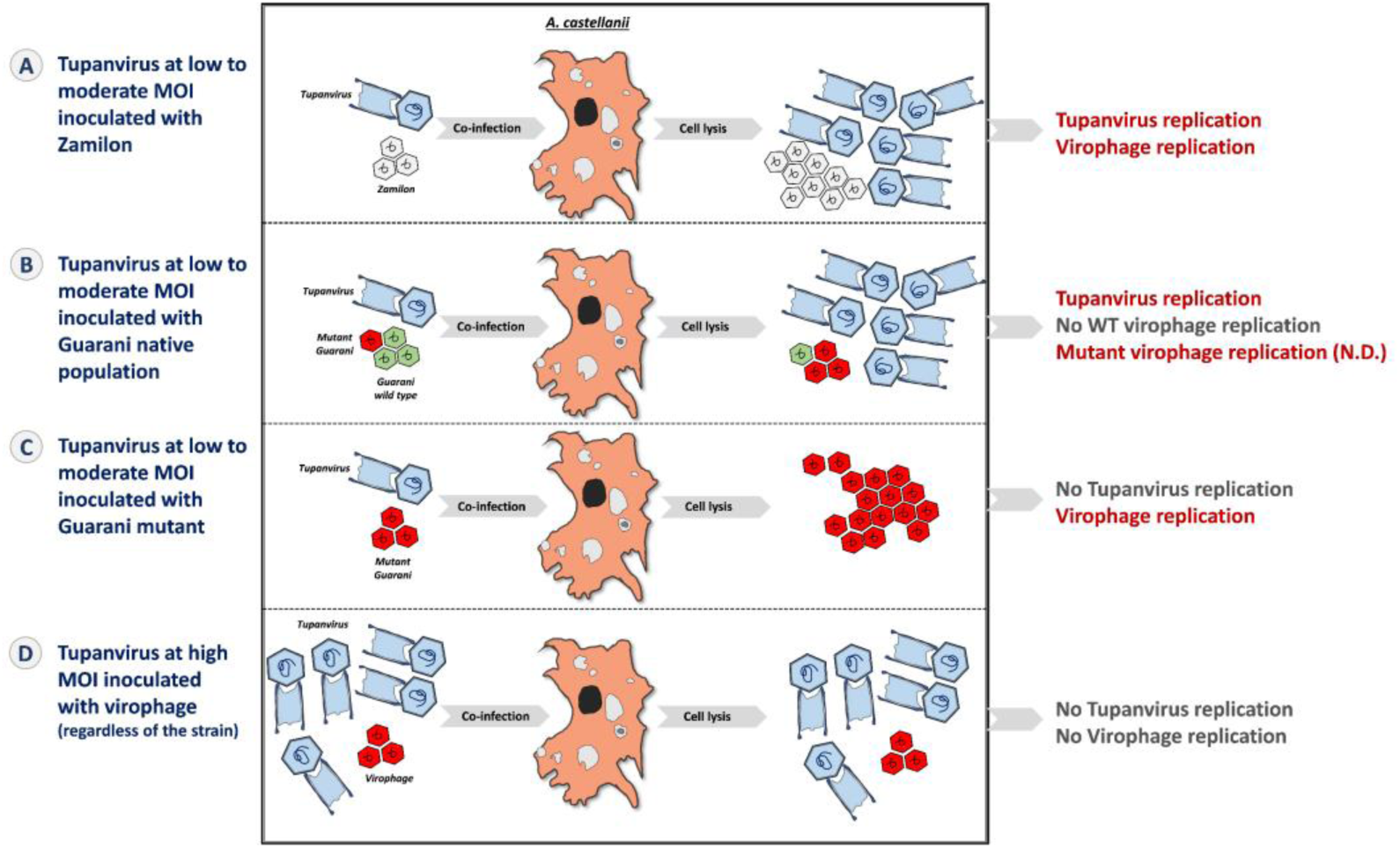
Scheme summarizing the different profiles of interaction between Tupanvirus and virophages described in our study. (A) Coinfection of *A. castellanii* with Zamilon and Tupanvirus at low to moderate MOI leads to replication of both Tupanvirus and virophage. (**B**). Guarani wild-type (WT) is not able to replicate with Tupanvirus in primo coinfection, but this first passage allows the selection of a mutant genotype adapted to the virus. The mutant Guarani is able to propagate with Tupanvirus and seems to be highly virulent, leading to the abolition of Tupanvirus morphogenesis. (**C**). Tupanvirus at high MOI triggers its cytotoxic profile in host cells. Although the virus cannot replicate at this MOI, this feature allows it to prevent virophage parasitism (**D**). N.D.: Not detected.

To the best of our knowledge, our study was the first to provide evidence of virophage abilities to expand their host range to infect new giant viruses. This study also highlighted a relevant impact of this host adaptation on giant virus and virophage replication and on lysis of their host cells. Thus, our results help to elucidate the parasitic lifestyle of virophages and their ecological influence on giant virus and protist populations.

## Materials and Methods

### 1. Tupanvirus production

*Acanthamoeba castellanii* cells (ATCC 30010) cultivated in PYG (Peptone, Yeast extract, Glucose) medium were used to produce Tupanvirus Deep Ocean and Tupanvirus Soda Lake. Suspensions containing 7.10^6^ cells plated in T175 flasks (Thermo Fisher Scientific, USA) were inoculated with each virus at an MOI of 0.02 and incubated at 30°C. After complete lysis of the cells, each virus supernatant was collected and centrifuged at 1000 g. The obtained supernatant was then filtered through a 0.8-μm membrane to remove amoeba debris. Each viral pellet was submitted to three cycles of wash with Page’s modified Neff’s amoeba saline (PAS) by ultracentrifugation at 14 000 g for 1 h. Finally, each virus was purified through ultracentrifugation across a 25% sucrose cushion at 14 000 g for 1 h.

### 2. Virophage production

Acanthamoeba polyphaga mimivirus (APMV) was used to propagate the Sputnik and Guarani virophages. To produce the Zamilon virophage, Megavirus Courdo 11 was used as a giant virus host. *A. castellanii* trophozoites at a concentration of 5.10^5^ cell/ml in PYG were inoculated with each virus at an MOI of 10 and each virophage. After lysis of the cells, the supernatant containing both virus and virophage particles was centrifuged at 10 000 g for 10 min and then filtered through 0.8-, 0.45- and 0.22-µm-pore filters to remove giant virus particles and residual amoebas. The virophage particles were concentrated by ultracentrifugation at 100 000 g for 2 h, and the pellet was resuspended with PAS. Each virophage was then purified through ultracentrifugation at 100 000 g for 2 h across a 15% sucrose cushion. A pure highly concentrated suspension was finally obtained for each virophage, in which the absence of mimivirus particles was confirmed by negative staining electron microscopy and inoculation in amoebas.

### 3. Host range studies

*A. castellanii* cells, the cellular support of the system, were resuspended three times in PAS. Ten milliliters of rinsed amoeba at 5×10^5^ cell/ml were simultaneously inoculated with Tupanvirus Deep Ocean or Tupanvirus Soda Lake and each virophage at MOIs of 10. The cocultures were incubated for 1 h at 30°C, and then extracellular giant virus and virophage particles were eliminated by 3 successive rounds of centrifugation and resuspension in PAS (1000 g for 10 min). The cocultures were then submitted to a second incubation at 30 °C. This time point was defined as H0. Each Tupanvirus strain was separately incubated with amoeba in the absence of virophage to serve as a negative control. At time points 0, 24 and 48 h p.i., a 200 µl aliquot of each coculture was collected for real-time PCR targeting the Major capsid protein (MCP) gene of each virophage (Table 1).

**Table 1:**
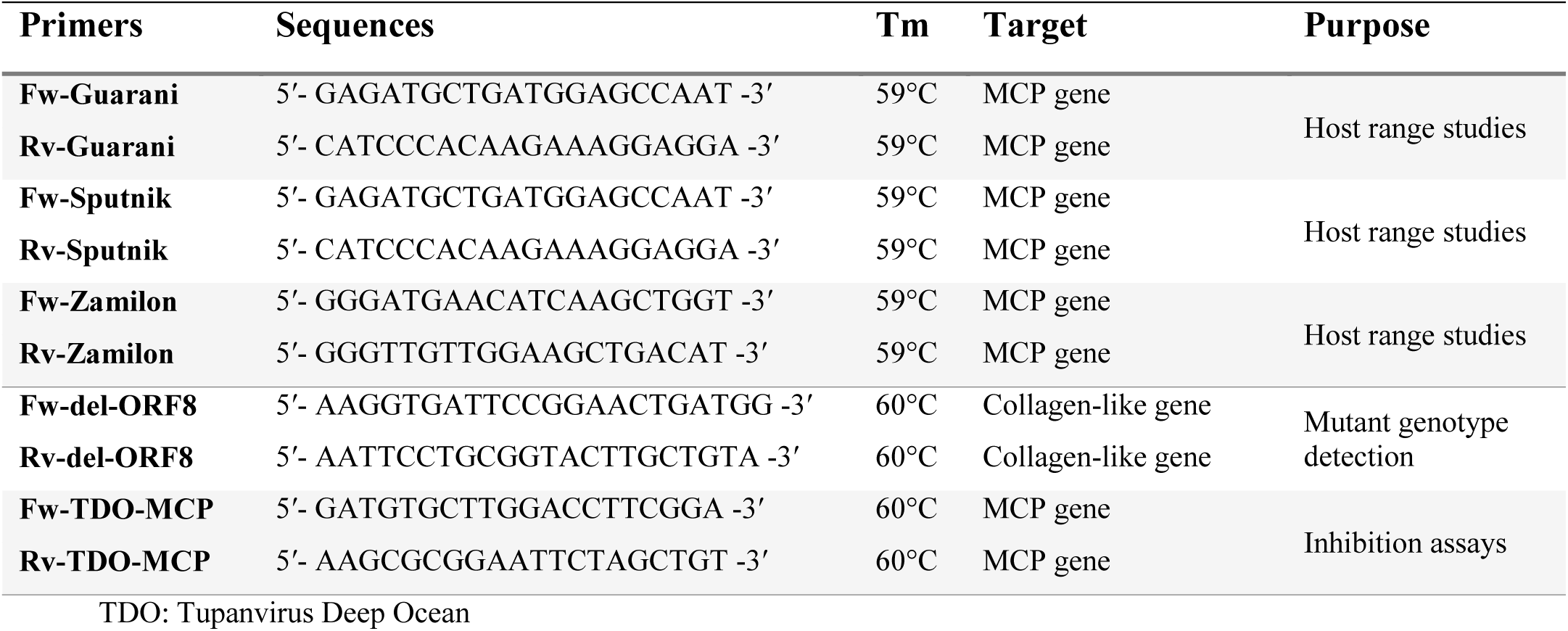
Primers used for host range studies, mutant Guarani detection and characterization.

To study the host range expansion of Guarani, the supernatant, obtained after lysis of the host cells coinfected with Guarani and Tupanvirus Deep Ocean or Tupanvirus Soda Lake, was filtrated through a 0.22-µm membrane to remove Tupanvirus particles. One hundred microliters of the filtrate containing only Guarani particles was subsequently used to infect fresh *A. castellanii* cells in PAS simultaneously inoculated with Tupanvirus Deep Ocean or Tupanvirus Soda Lake at MOIs of 10. The replication of the virophage was then quantified as described above. The experiment was carried out three times independently in duplicate.

## 4. Real-time PCR

DNA extraction and PCRs were performed as described by Mougari et al^24^. All the primers used here are listed in Table 1. Virophage replication with each Tupanvirus strain was calculated by the ΔCt method, considering the difference between times 0 and 48 h p.i.

## 5. Detection and characterization of the mutant genotype

### 5.1. Selection of the mutant Guarani

Two T175 flasks (Thermo Fisher Scientific, USA) containing each of 20 million *A. castellanii* cells in PYG medium were inoculated with Tupanvirus Deep Ocean and Guarani wild-type (propagated with APMV) at MOIs of 10. The coculture was incubated at 30 °C. After complete lysis of the cells, the virus-virophage supernatant collected from each flask was used to infect 10 more T175 flasks, and the cocultures were incubated at 30 °C. Approximatively 1 L of the virus-virophage supernatant was collected from all cocultures. The virophage particles were then purified as described above and subsequently submitted to genome sequencing. The same procedure was repeated for Tupanvirus Soda Lake and APMV (control).

### 5.2. Genome sequencing and analyses

To investigate whether the Guarani obtained from Tupanvirus Deep Ocean and Tupanvirus Soda Lake were genetically different from that propagated with APMV, the virophage cultivated with each virus was submitted to genome sequencing with Illumina MiSeq. The genome of each virophage was assembled and analyzed as previously described by Mougari et al^24^. Comparative genomic analysis and genome alignment were then conducted using Muscle software and BLASTn alignment (Basic Local Alignment Search Tool)^56^.

### 5.3. Standard PCR and Sanger sequencing

A specific PCR system targeting the collagen-like protein (ORF8) was designed and performed on the DNA extracted from supernatant containing the mutant genotype to confirm the occurrence of deletion of the 81 nucleotide-long sequence in this strain (Fig. 3a). The details of the primers used are listed in Table 1, and the PCR product was visualized on an agarose 2% gel using SYBR safe buffer (Invitrogen, USA). The Qiagen gel extraction kit was used to recover the band corresponding to each genotype of Guarani detected in Tupanvirus supernatant (wild-type and mutant) from the 2% agarose gel according to the manufacturer’s instructions. Sanger sequencing analyses were then performed in an Applied Biosystems^®^ 3130/3130xl Genetic Analyzer (Thermo Fisher, USA) using a Big Dye Terminator Cycle Sequencing Kit (Thermo Fisher, USA) according to the manufacturer’s instructions. The obtained sequences were assembled with ChromasPro 1.7.7 software (Technelysium Pty Ltd, Australia).

## 6. Passage experiments and inhibition assays

To investigate whether mutant Guarani was able to maintain the deletion mutation during several passages with Tupanvirus, *A. castellanii* cells at a concentration of 5.10^5^ cell/ml in 10 ml PYG medium were inoculated with Tupanvirus Deep Ocean and Guarani wild-type genotype at MOIs of 10. The coculture was incubated at 30°C until complete lysis of amoebas was observed. To perform serial flask passaging, 100 µl of the virus-virophage supernatant collected from this primo coculture was used to infect fresh *A. castellanii* cells seeded at a 5. 10^5^ cells/ml density in 10 ml of PYG medium. The procedure was repeated 5 times. After each passage, 200 µl aliquots of the coculture were collected for PCR targeting the deletion site in Guarani.

To evaluate whether mutant Guarani was able to inhibit the production of Tupanvirus infectious particles during a passage experiment, *A. castellanii* cells at a 5. 10^5^ cells/ml density in 10 ml PYG medium were simultaneously coinfected with Tupanvirus Deep Ocean at an MOI of 10 and 100 µl of Guarani mixture containing both wild-type and mutant genotypes (according to PCR). The Guarani used here was isolated from the second passage with Tupanvirus where the mutation was detected. The serial passaging experiment was performed as described above. After each passage, the titer of infectious particles was quantified by end point dilution.

To study the effect of mutant Guarani on the DNA replication and virion production of Tupanvirus during a one-step growth curve, cells were coinfected with Tupanvirus Deep Ocean and mutant Guarani at MOIs of 10. The Guarani used in this experiment was isolated and then purified from the second passage with Tupanvirus. To assess the effect on DNA replication, a 200-µl aliquot of the coculture was collected after 0 and 48 h p.i. and submitted to DNA extraction and then to real-time PCR that targets the MCP encoding gene in the Tupanvirus Deep Ocean (Table 1). To evaluate the effect on the production of Tupanvirus virions, a 1 ml aliquot was collected at times 0, 6, 12, 16, 24, 32, 48 and 72 h p.i. The virus supernatant was frozen and thawed three times to release the virions, and the titer was then determined at each time point by end point dilution.

To perform the dose-response to virophage, the same procedure was repeated using different MOIs of the giant virus (Tupanvirus or APMV) and the virophage (Guarani or Zamilon). The supernatant was then collected for end point dilution after complete lysis of cells or 5 days p.i. in the absence of cell lysis.

## 7. Virus titration

Virus titration was performed as previously described^24^. To inhibit the virophage and avoid interference with giant virus multiplication during the end point dilution method, each virus supernatant was heated for 30 min at 55°C. This treatment allowed us to inactivate the virophage without a significant decrease in the virus titer.

## 8. Transmission electron microscopy (TEM)

TEM experiments were performed as previously described by Mougari et al^24^

## 9. Ribosomal RNA shutdown and virophage replications assays

To investigate whether the cytotoxic profile of Tupanvirus associated with ribosomal RNA (rRNA) shutdown was able to prevent virophage replication with the virus and the capacity of mutant Guarani to inhibit this phenotype, 1 million *A. castellanii* cells were infected with Tupanvirus Deep Ocean or Moumouvirus at MOIs of 100 and each virophage at MOIs of 10. The evaluation of the rRNA shutdown for each condition was conducted as previously described by Abrahao et al^20^. The DNA replication of virophages was quantified as detailed above.

## Supplementary data

**Table S1: Nucleotide sequences of the collagen-like gene in wild-type and mutant Guarani.**

**Supplementary Figure 1: Location of the deletion in the collagen-like gene of Guarani.** The collagen-like repeats and their amino acid motifs are highlighted in colors. Only a part of the gene is represented here.

**Supplementary Figure 2: Selective inactivation of the mutant virophage using heat treatment at 55 °C for 30 min.** (**a**) The supernatant collected from passage 2 with Tupanvirus was filtered through a 0.22-µm-pore filter to remove giant virus particles. The obtained supernatant was then serially diluted from 10^-1^ to 10^-5^. Each dilution was then submitted to heat treatment and subsequently used to coinfect *A. castellanii* cells simultaneously inoculated with Tupanvirus Deep Ocean. The virophage did not replicate in all conditions, indicating that heat treatment successively inactivated it. (**b**) Different concentrations of Tupanvirus Deep Ocean were submitted to heat treatment and then titrated using end point dilution. We did not observe any change in the titer of viable particles after the treatment. (**c**) *A. castellanii* cells were inoculated with Tupanvirus at an MOI of 1 and mutant Guarani at an MOI of 10. Prior to coinfection, virus-virophage suspensions were mixed, incubated to allow composite formation and then submitted to heat treatment. After lysis, the titer of infectious particles was measured by end point dilution. Heat treatment completely prevented virophage inhibition and allowed Tupanvirus to replicate.

**Supplementary Figure 3: Simultaneous production of Tupanvirus and Zamilon virions in coinfected *A. castellanii* cells.** (**a** and **b**) *A. castellanii* cells were simultaneously inoculated with Tupanvirus Deep Ocean and Zamilon at MOIs of 10. Zamilon virophage progeny (white arrows), Tupanvirus particles (red arrows).

**Supplementary Figure 4: Detection of deletion of the 81-nucleotide sequence in Sputnik virophage by PCR.** To detect the mutation in Sputnik, the virophage was propagated with APMV and purified as described above. Sputnik DNA was extracted from the purified virophage. Five nanograms of Sputnik DNA were then used as a template for PCR. Sanger sequencing on each band confirmed the mutation. The primers used for PCR and Sanger sequencing are the same as Guarani.

**Supplementary Figure 5: Effect of conservation on mutant Guarani virion viability.** The virophage was isolated from Tupanvirus supernatant (passage 2) and then incubated at room temperature for 2 months or conserved at 4°C for 5 months, approximatively. Virophage ability to replicate with Tupanvirus was then assessed by real-time PCR as described above. The virophage did not replicate in either condition. The virophage survived other incubation times (15 days and 1 month at room temperature and 2 months at 4°C, data not shown).

## Funding

This work was supported by a grant from the French State managed by the National Research Agency under the “Investissements d’avenir” (Investments for the Future) program with the reference ANR-10-IAHU-03 (Méditerranée Infection) and by Région Provence-Alpes-Côte-d’Azur and European funding FEDER PRIMMI. Said Mougari was supported by the Foundation Méditerranée Infection scholarship.

## Author Contributions

**Said Mougari** conceived the study, designed and performed the experiments, collected, analyzed and interpreted the data, drafted the text and wrote the manuscript. **Nisrine Chelkha** and **Said Mougari** analyzed genomic data. **Dehia Sahmi-Bounsiar** performed experiments. **Fabrizio Di Pinto** and **Said Mougari** performed electronic microscopy experiments. **Colson Philippe** analyzed genomic data, reviewed the manuscript and contributed to the final version. **Bernard La Scola** and **Jonatas Abrahao** conceived the study, designed the experiments and wrote the manuscript.

## Competing interests

The authors declare no competing financial interests.

