## Supplementary material for "First evidence of host range expansion in virophages and its potential impact on giant viruses and host cells"

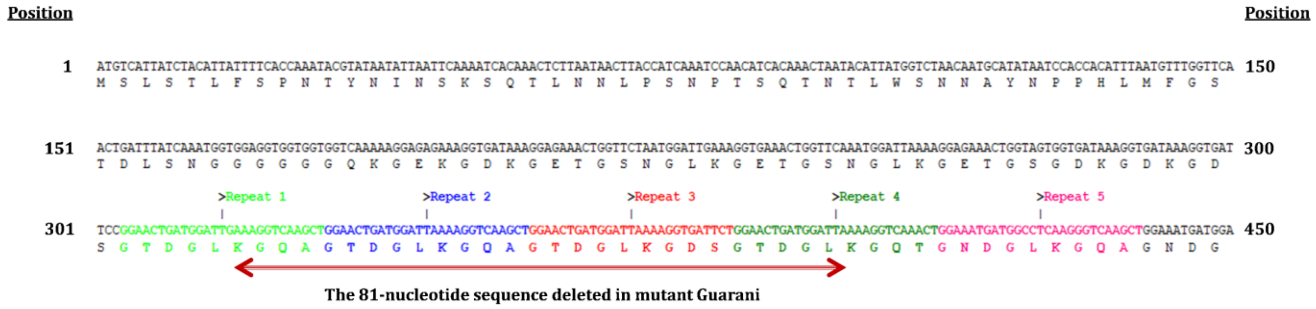


### Supplementary Figure 1

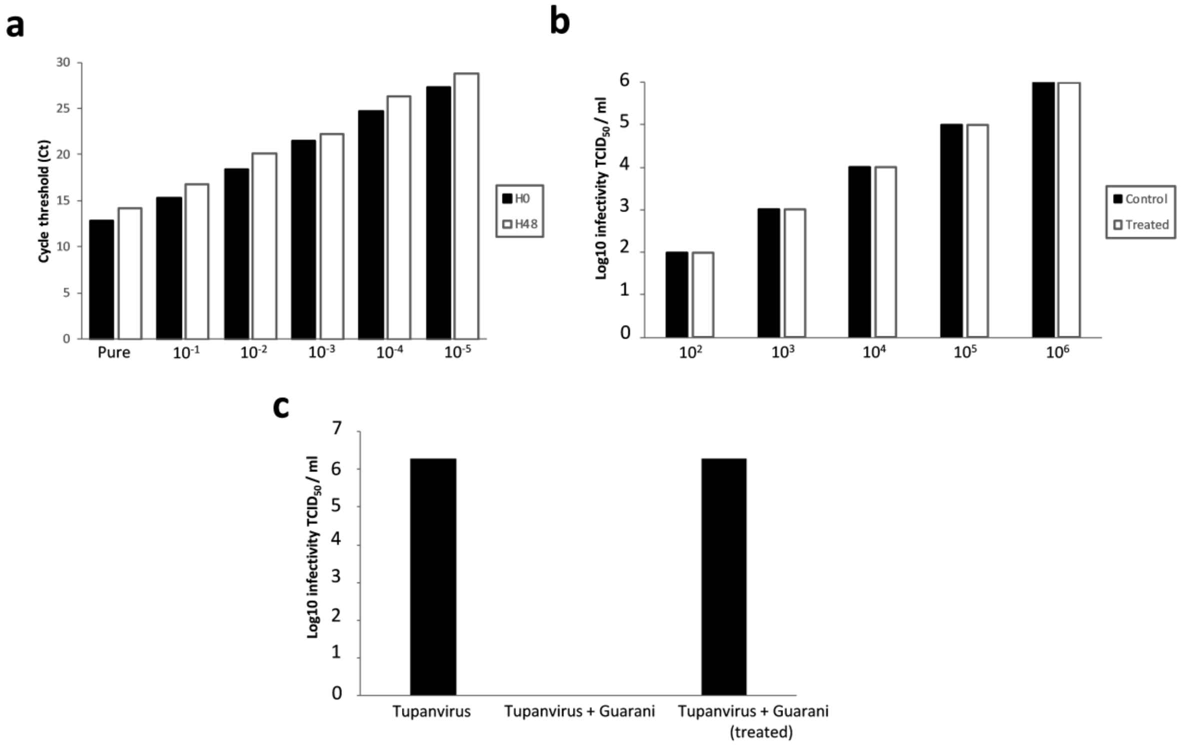


**Supplementary Figure 2**

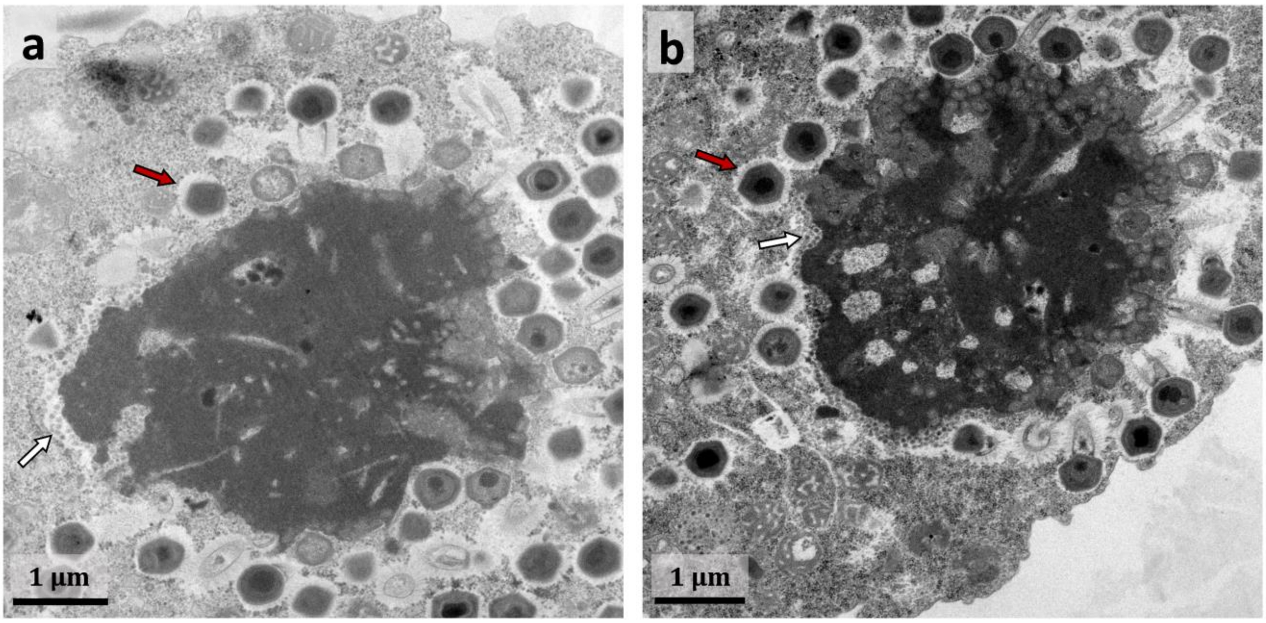


**Supplementary Figure 3**

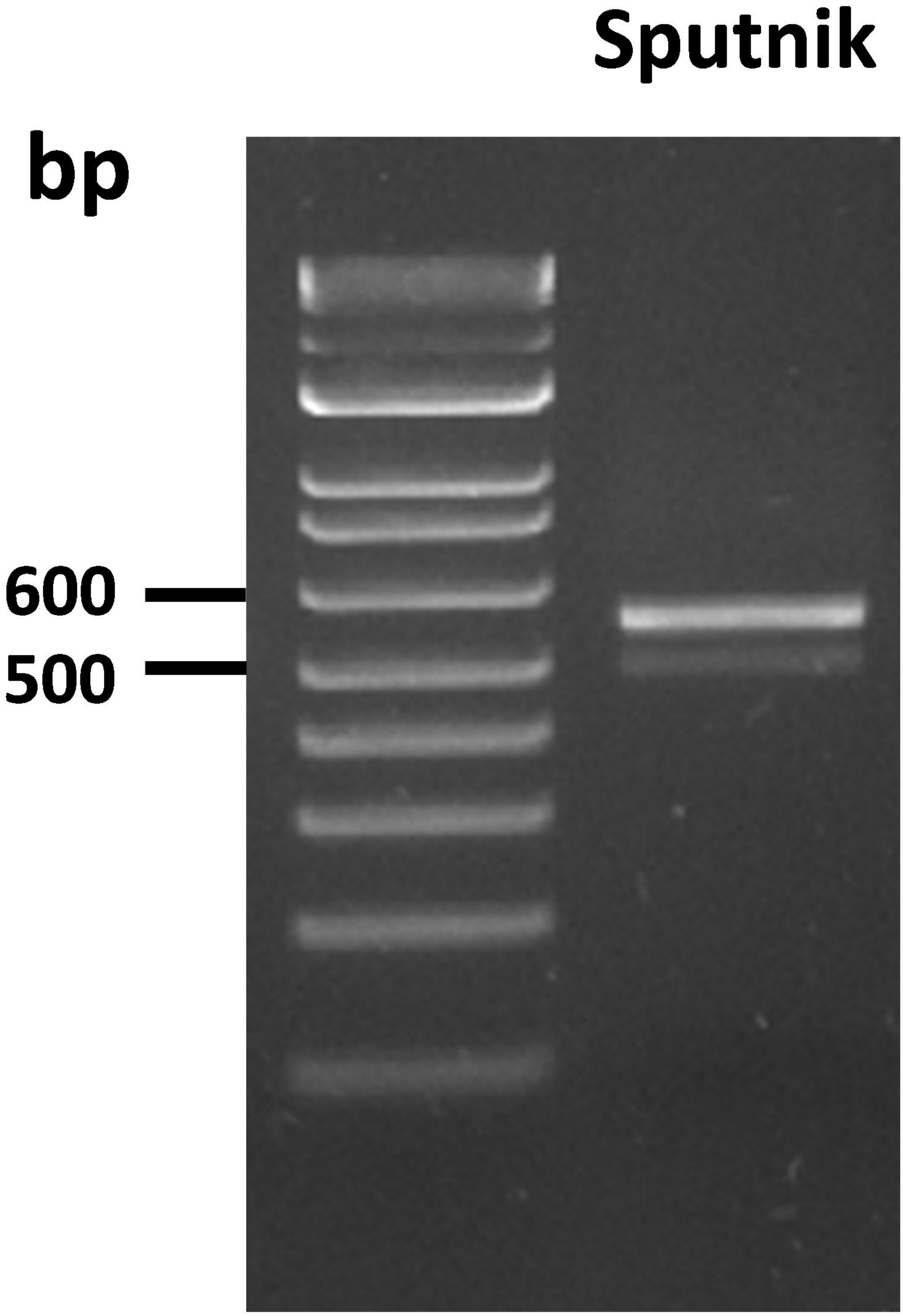


**Supplementary Figure 4**

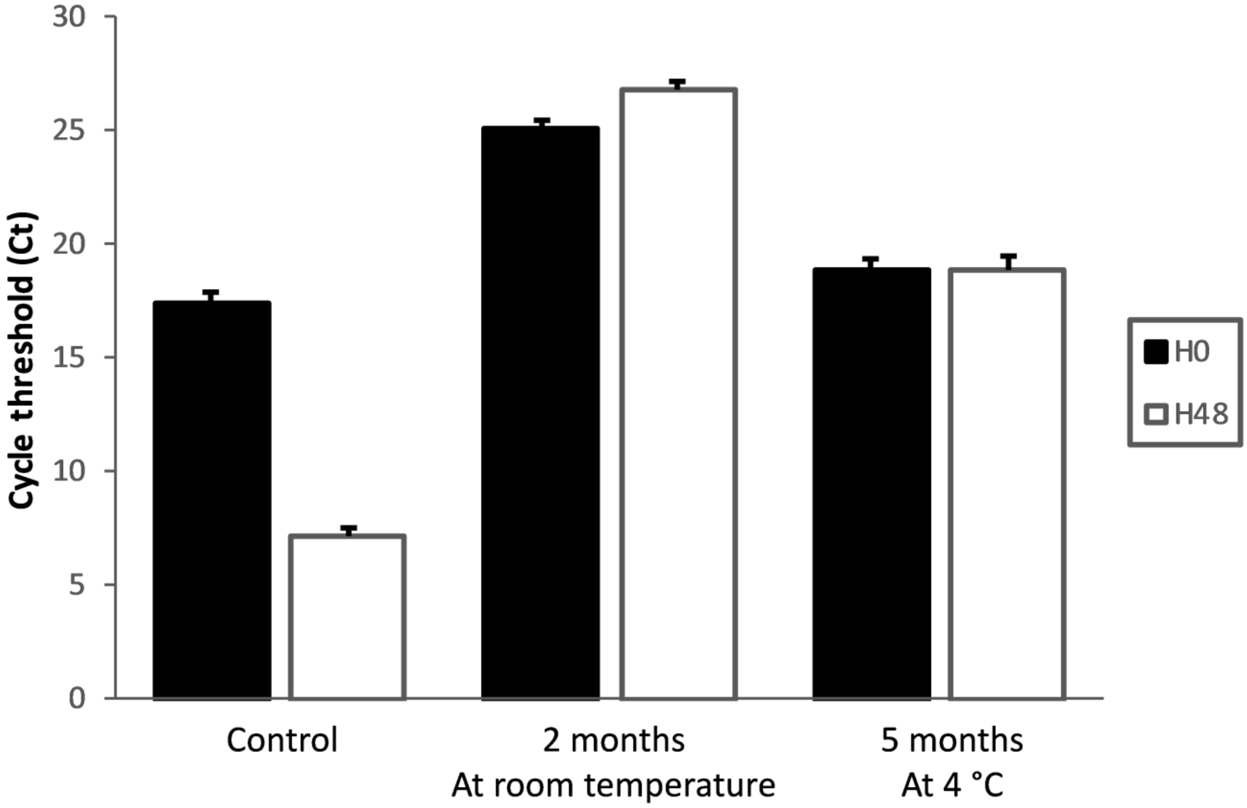


**Supplementary Figure 5**
